# The representation of vocalizations in structured noise in the mouse auditory cortex

**DOI:** 10.64898/2026.09.28.753104

**Authors:** Z. Cai, J. Peters, S. Arends, B. Englitz

## Abstract

Efficient processing of complex acoustic signals within noisy environments is a fundamental challenge for the auditory system. This study investigates the cortical representation of structured noise and vocalizations embedded in it using widefield calcium imaging with the high-speed Calcium indicator jGCaMP8m in the mouse auditory cortex. The spectrotemporal statistics of the structured noise were controllable, including its temporal variance and cross-frequency correlations.

We find that structured noise invokes spatially and temporally complex neural responses across the auditory cortex that reflect the spectrotemporal properties of the noise. We identified partially distinct noise response components: noise onset responses showed stronger overlap with primary than secondary areas, while later suppression components did not show a significant primary-secondary bias. Vocalization-evoked responses were significantly attenuated in the presence of noise compared to silent contexts, exhibiting reduced onset magnitude and altered adaptation dynamics. Using decoding we find the neural representation of vocalizations to become more discriminable with longer preceding noise exposure. Furthermore, structured noise cochleograms could be reconstructed from neural population activity, with performance depending on stimulus statistics, tending to be higher in primary than secondary auditory cortex while remaining substantial across both.

These findings suggest that the mouse auditory cortex represents acoustic targets in noise through context-dependent and spatially distributed population activity, shaped by both the statistical structure and temporal history of the acoustic background.

**Highlights:**

- Structured noise invokes a spatially and temporally complex neural response across auditory cortex
- Spectrotemporal properties of the noise can be reconstructed from the spatiotemporal response
- Decoding quality for vocalizations improves with exposure to the noise during passive listening
- Acoustic information is distributed across auditory cortex rather than confined to a single subdivision

## Introduction

Natural listening rarely involves isolated sounds. Behaviorally relevant signals, such as vocalizations, are usually embedded in complex acoustic backgrounds produced by other animals, movement, wind, water, or other environmental sources. Because these backgrounds are dynamic and statistically structured, perceiving a target sound requires the auditory system to extract relevant acoustic events from a continuously changing context, a challenge often discussed in relation to auditory scene analysis and the cocktail-party problem (Bregman, 1990; Cherry, 1953; Middlebrooks et al., 2017). This requires more than detecting sound energy at a particular frequency or time: the same target sound may be easy or difficult to detect depending on the structure of the surrounding background. Understanding how the brain represents sounds in naturalistic environments, or more generally in statistically structured acoustic contexts, therefore requires considering not only the target sound itself, but also the recent acoustic context in which it occurs (Theunissen & Elie, 2014; Willmore & King, 2023).

One way the auditory system may solve this problem is by representing and adapting to the statistical structure of sound. Natural sounds contain regularities in their spectrotemporal modulations, and auditory neurons are sensitive to these structures rather than simply encoding isolated acoustic features (Escabí & Read, 2003; Singh & Theunissen, 2003; Theunissen & Elie, 2014). Such sensitivity provides a basis for estimating the recent acoustic environment. Once the statistics of the background are estimated, neural responses can adapt to them, for example through changes in gain, response magnitude, or sensitivity to contrast (Cooke et al., 2018; Rabinowitz et al., 2011, 2012; Willmore & King, 2023). This adaptation may suppress predictable or redundant components of the background while preserving responses to acoustic events that deviate from it (Barlow, 1961). In this view, target-in-noise perception is not only a problem of detecting a signal against masking energy, but also a problem of using recent sound statistics to define what is expected and what is informative.

To study this process experimentally, the acoustic background must be complex enough to contain meaningful statistical structure, but controlled enough to isolate which statistics matter. Sound textures provide one such framework. In texture models, naturalistic background sounds can be described by summary statistics of auditory-filter outputs, including the variability of activity within frequency channels and the correlation of amplitude fluctuations across channels (Hicks & McDermott, 2024; McDermott et al., 2013; McDermott & Simoncelli, 2011; McWalter & McDermott, 2018). The distribution of these statistical features has also been characterized across natural sound textures, and neural responses in the auditory midbrain and cortex are sensitive to such texture statistics (Mishra et al., 2021; Peng et al., 2024). This makes it possible to synthesize artificial sounds that are not natural recordings, but nevertheless contain controlled forms of naturalistic structure. Similarly, spectrotemporally modulated stimuli such as ripples and temporally orthogonal ripple combinations (TORCs) provide a way to probe auditory responses with controlled modulation patterns over time and frequency (Depireux et al., 2001; Klein et al., 2000). Together, these approaches provide an intermediate strategy between simple tones or white noise, which lack naturalistic structure, and natural recordings, in which many acoustic features covary. They therefore allow specific statistical features of sound to be manipulated while retaining structured acoustic complexity that captures certain aspects of natural sounds.

To understand how structured backgrounds shape the cortical representation of behaviorally relevant sounds such as vocalizations, it is important to measure activity across the auditory cortex while preserving temporal detail. Complex sounds can recruit multiple auditory fields and evoke response components on different timescales. While electrophysiological recordings provide high temporal resolution (Alishbayli et al., 2025), they cannot comprehensively sample activity across the entire auditory cortex. Widefield calcium imaging provides such spatial coverage (Issa et al., 2014; Romero et al., 2020). And recent rapid calcium indicators enable these large-scale responses to be resolved with improved temporal fidelity (Zhang et al., 2023; Peters et al., 2026).

Here, we used widefield imaging with jGCaMP8m to record activity across the auditory cortex during presentation of structured noise containing randomly occurring mouse vocalizations. We examined how the statistical structure and preceding duration of the acoustic background influenced responses to both the noise and embedded vocalizations. We further used population decoding and stimulus reconstruction to assess how information about these sounds was represented across cortical activity. We find that neural responses depend on both the statistics and temporal history of the background, and that information about the acoustic context and embedded vocalizations is represented across distributed cortical population activity.

## Results

We investigated neural responses from the auditory cortex in normal hearing CBA/JRj mice (N = 7) to conspecific vocalizations. These were presented in silence or embedded in structured noise with a range of temporal variances and spectral correlations. We recorded from the entire auditory cortex using widefield imaging while the mice listened passively (Fig. 1A) after multiple injections of the fast and bright calcium indicator jGCaMP8m during the implant surgery (Fig. 1B).

**Figure 1:**
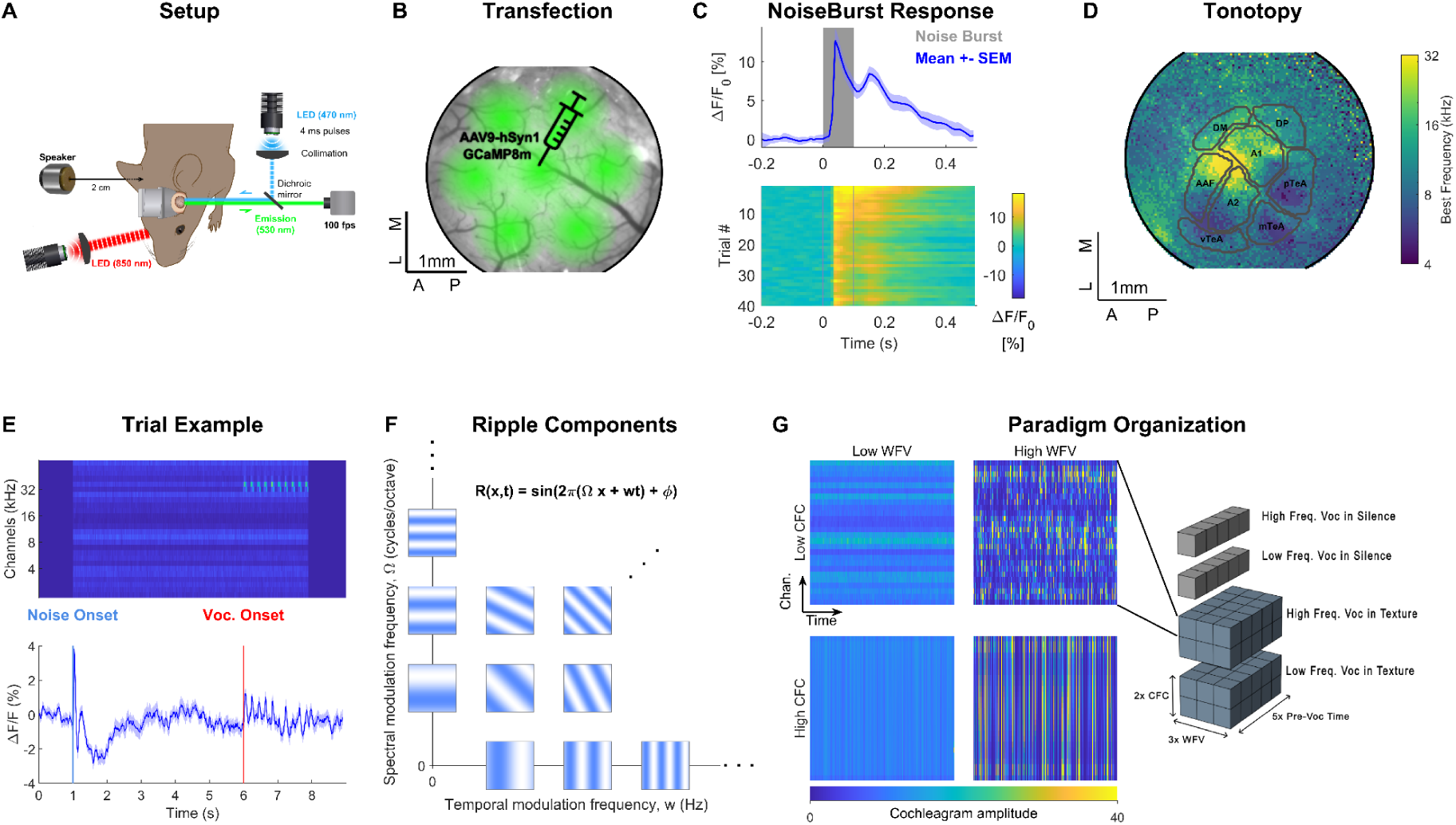
Experimental and stimulus design. **A** Neural activity in the auditory cortex (AC) was imaged through a cranial window using the fast and bright jGCaMP8m indicator using widefield imaging. Excitation light (470 nm) was delivered through the objective and emitted fluorescence light (525 nm) was recorded at 100 fps. Acoustic stimuli were delivered from a calibrated speaker positioned ∼2 cm from the contralateral ear. **B** To image the entire auditory cortex rather uniformly, transfection with the calcium indicator was performed in multiple, hexagonally-arranged injection sites. The fluorescence image shows broad expression across auditory cortex, with brighter regions corresponding to injection centers. **C** Mean fluorescence response (ΔF/F, mean ± SEM) from an ROI in area A1 to a broadband pink noise (grey area, 100 ms, 40 repetitions, intertrial interval: 4-5 s). Bottom: single-trial responses. **D** Fields of AC were defined based on latency and frequency response maps (’Tonotopic map’), computed from responses to pure tones. Boundaries of approximate cortical fields are overlaid on the tonotopic map (AAF: Anterior Auditory Field, DM: Dorsomedial Field, DP: dorsoposterior field, v/m/pTeA: ventral/medial/posterior temporal association area). Derived from a representative animal’s tonotopic mapping. **E** Acoustic stimuli were vocalizations embedded in structured noise stimuli. Vocalizations were randomly placed in time relative to noise onset (0.5/1/3/5/7 s) and frequency (8/32 kHz), and adjusted in intensity to be 5 dB above the noise level in their frequency bins. Sample stimulus shows a cochleogram of the stimulus with the vocalizations at 32 kHz, repeated 10 times. The neural response showed a highly dynamic temporal evolution. **F** Structured noise was created using sums of spectrotemporal ripples to limit the modulations to the range represented in AC, called TORCs (Klein et al. 2000). Ripples were defined by their temporal modulation frequency (w) and spectral modulation frequency (Ω). **G** TORCs were modified to differ in their cross-frequency correlations (CFC, top vs. bottom) and degree of within frequency channel variance (WFV, left vs. right). Sample spectrograms were computed as cochleograms. Overall (right), this led to 70 different stimuli (5 pre-vocalization times, 3 WFV, 2 CFC, 2 vocalization frequencies), including a condition where vocalizations were presented in silence (5 pre-vocalization times, 2 vocalization frequencies).

### Auditory cortex exhibits reliable, structured, and stimulus-dependent responses

We first characterized the neural responses in the auditory cortex using basic sounds to estimate a reference map of cortical subareas per animal. Generally, response fidelity was high, showing rapid and precisely timed responses even on single trials (Fig. 1C, broadband pink noise) in accordance with the fast dynamics and high sensitivity of jGCaMP8m (***τ***_onset_ = ∼5 ms, ***τ***_recovery_ = 70 ms, including internal Ca-dynamics).

To characterize the spatial organization of these responses, we computed tonotopic maps based on pure tone stimulation (Fig. 1D). Distinct auditory fields were identified according to best frequency and response latency (Fig. 1D), providing a functional reference map for subsequent analyses. Specifically we first coarsely distinguished primary and secondary areas based on response latency (Peters et al., 2026), and then subdivided them into the areas A1/A2/AAF, and TeA/DP/DM in accordance with previous studies (Issa et al., 2014; Romero et al., 2020, see Methods for details).

Next, we presented composite sounds in which a sequence of vocalizations was embedded inside a complex acoustic context, which we refer to as structured noise (Fig. 1E top). These composite sounds model the challenge of a listening task in which a target sound (e.g. speech) occurs in the midst of an environmental sound. Importantly, in each trial the vocalizations are positioned at a randomly drawn time-point ([0.5,1,3,5,7] s) and frequency ({8,32} kHz), preventing the animal from developing a basic time or frequency-based listening strategy. Further, in each trial the statistics of the noise are randomly drawn from a limited set (see below). Therefore, a listener could optimize performance on this task by adapting to the statistics inside a given trial, which could benefit from longer exposure to the sound (Alishbayli et al., 2025). The structured noise was based on a sum of spectrotemporal ripples, which limits the modulations to a range that can be represented in the cortex (Fig. 1F, Depireux et al., 2001; Klein et al., 2000). The noise properties were selected to modulate the correlations across frequencies (termed Cross-frequency correlations, CFC) and the variance within a frequency band (termed within-frequency variance, WFV). These properties provide additional constraints or variability, respectively, and should therefore have an influence on quality with which the noise statistics can be estimated and/or the vocalization can be detected against the noise. Overall, this resulted in 70 stimulus conditions (5 onset times X 3 WFV X 2 CFC X 2 vocalization frequencies + 5 onset times X 2 vocalization frequencies in silence, see Fig. 1G).

When exposed to structured noise and vocalization stimuli, neural responses showed rich temporal dynamics that reflected both ongoing noise stimuli and transient acoustic events (Fig. 1E). These responses evolved over time, indicating that auditory cortical activity integrates stimulus history and instantaneous input.

*Structured noise reveals distinct spatial and temporal components of cortical responses Spatial organization of onset and suppression response components*: To capture different aspects of the response to the structured noise, we defined two per-pixel cortical maps we considered relevant indicators: (1) Noise Onset Map (NOM): The positive peak of the onset response to the structured noise (Fig. 2A-F, orange) was computed shortly after stimulus onset (0-70 ms) and was considered as the parts of the auditory cortex that responded most strongly to the noise. (2) Noise Minimum Map (NMM): This was defined based on the response minimum after the onset (Fig. 2A-F, green) and was considered to indicate the parts of the auditory cortex that show the strongest suppression, typically below baseline. Both maps were computed separately for each animal and stimulus condition (Fig. 2A-F show a representative animal).

**Figure 2:**
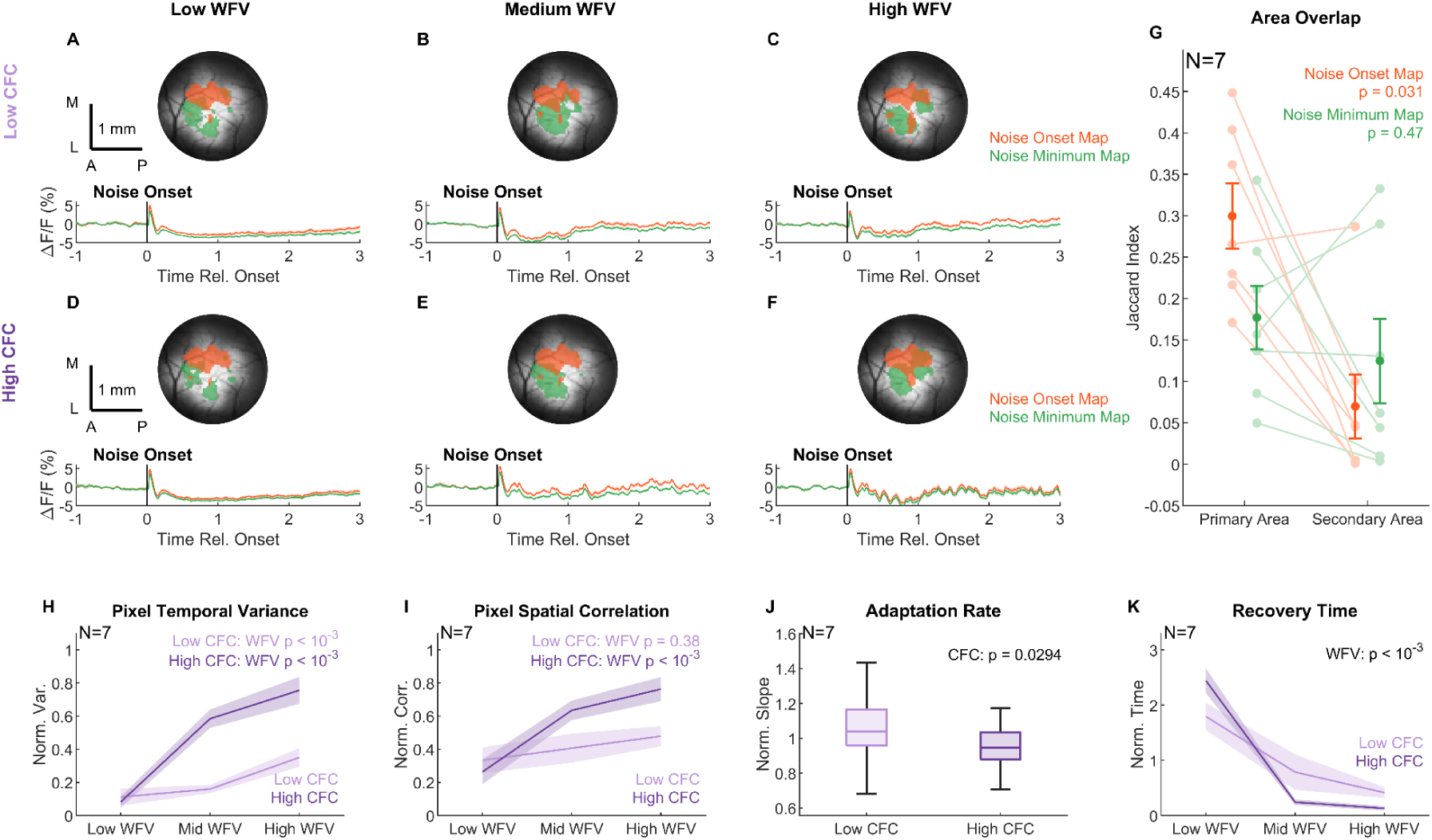
Spatial organization of cortical activity to noise onset and temporal evolution A–F. Neural responses of a representative animal to structured noise stimuli with different spectrotemporal statistics, varying in cross-frequency correlation (CFC, top vs bottom) and within-frequency variance (WFV, left to right). For each panel, response maps are overlaid on average fluorescence images (top), and mean traces extracted from the response-defined areas are shown (bottom). Response maps highlight areas with the largest onset responses to the noise (orange), defined from the per-pixel maximum ΔF/F during an early post-onset period, and areas showing the strongest suppression (green), defined from the lowest 5% of ΔF/F values during a later adaptation period. Binary response maps were obtained by combining response maps across pre-vocalization times and applying a z-score threshold (z > 1.5); the same maps were used for the analysis in G. **G** Quantification of map overlap with primary and secondary auditory regions. Each line connects values from the same animal. The noise onset map showed significantly (p = 0.031, WSRT paired comparisons between primary and secondary areas within each map type) greater overlap with primary than secondary auditory areas, whereas the noise minimum map showed no significant difference between primary and secondary areas. Overlap quantified as Jaccard index between maps and auditory fields. **H** Pixel-wise temporal variance during structured-noise responses. Variance was computed across temporal bins for each pixel and averaged within the craniotomy mask (normalized within animal). Increasing WFV increased cortical response variability under both CFC conditions, with stronger increases and clear separation under high CFC at medium and high WFV. **I** Local spatial correlation during structured-noise responses. Correlation was computed between each pixel and nearby pixels and averaged within the craniotomy mask (normalized within animal). Increasing WFV increased local cortical correlation under high CFC (p < 0.001, LME, N = 7) but not significantly under low CFC (p = 0.38, LME, N = 7), resulting in greater local coordination for high-CFC than low-CFC textures, particularly at medium and high WFV. **J** The speed of adaptation after the onset, quantified as the slope between onset peak and response minimum, i.e. (R_onset − R_min) / (t_min − t_onset). Adaptation was stronger for structured noise with low CFC than with high CFC (p = 0.029, LME, N = 7). Values normalized within-animal by dividing by the animal’s mean value across conditions. **K** The time required to recover from adaptation was significantly shorter in higher WFV conditions, measured to 80% recovery of the strongest adaptation (see Methods). Values normalized within-animal as above. Statistical analysis via a linear mixed-effects model with animal as random effect (N = 7).

Mean response traces extracted from the onset-and minimum-defined areas exhibited quite similar temporal profiles, reflecting shared underlying population dynamics (Fig. 2A–F, bottom panels). Despite this similarity, clear differences in response magnitude were observed, with both maps emphasizing distinct components of the response.

Although the resulting areas were spatially overlapping, they showed distinct anatomical biases. The onset response map overlapped significantly more strongly with primary than secondary auditory cortex (Jaccard index 0.31 ± 0.04 vs 0.07 ± 0.04; p = 0.031, Wilcoxon signed-rank test (WSRT); Fig. 2G). In contrast, the minimum response map showed comparable overlap with primary and secondary auditory regions (p = 0.47, WSRT), rather than a significant bias toward either subdivision. This spatial organization suggests that early transient components are preferentially weighted toward primary auditory cortex, whereas later suppressive components are less restricted to primary fields and are more broadly distributed across the auditory cortex.

*Stimulus statistics shape cortical response dynamics*: Response traces varied systematically with stimulus statistics. Increasing WFV and CFC led to larger and more coherent fluctuations in activity (Fig. 2A–F, bottom panels), with high CFC and high WFV stimuli producing larger fluctuations over time, consistent with increased shared modulation across the population.

To further characterize texture-dependent cortical dynamics, we quantified pixel-wise temporal variance and local spatial correlation during structured-noise responses (Fig. 2H–I). Increasing WFV led to progressively larger temporal fluctuations in cortical activity under both CFC conditions (Fig. 2H). Notably, responses under high CFC became increasingly separated from low CFC at medium and high WFV, indicating stronger modulation of cortical activity. Similarly, local spatial correlation between nearby pixels increased with WFV for high-CFC textures (p < 0.001, linear mixed-effects (LME) model with WFV as a fixed effect and animal as a random effect, Fig. 2I), but not significantly for low-CFC textures (p = 0.38, LME, Fig. 2I). As a result, high-CFC textures exhibited greater local coordination than low-CFC textures, particularly at medium and high WFV. These changes in ongoing activity organization paralleled the stronger adaptation and altered recovery dynamics observed for highly structured textures.

These qualitative differences were accompanied by systematic changes in the temporal dynamics of the response. The rate of adaptation, quantified as the slope between the onset peak and subsequent minimum, was significantly modulated by CFC, with stronger adaptation observed for low CFC stimuli compared to high CFC (Fig. 2J, p=0.029, LME, N = 7). In contrast, recovery from suppression was primarily influenced by WFV: higher WFV conditions led to faster recovery toward baseline, whereas low WFV stimuli produced more prolonged suppression (Fig. 2K).

These results suggest a dissociation in how different statistical features of the stimulus shape cortical dynamics, with cross-frequency structure shaping adaptation strength and within-frequency variability governing recovery timescales.

### Vocalization responses across sensory contexts

We compared cortical responses to identical vocalizations presented either in silence or embedded in ongoing noise. Similar to noise response related maps, the corresponding cortical responsive regions were defined on a per-pixel basis by thresholding stimulus-evoked activity maps (see Methods). Both conditions activated largely overlapping cortical regions (Fig. 3A, red and blue), indicating that vocalizations are represented in similar cortical locations across contexts.

**Figure 3.**
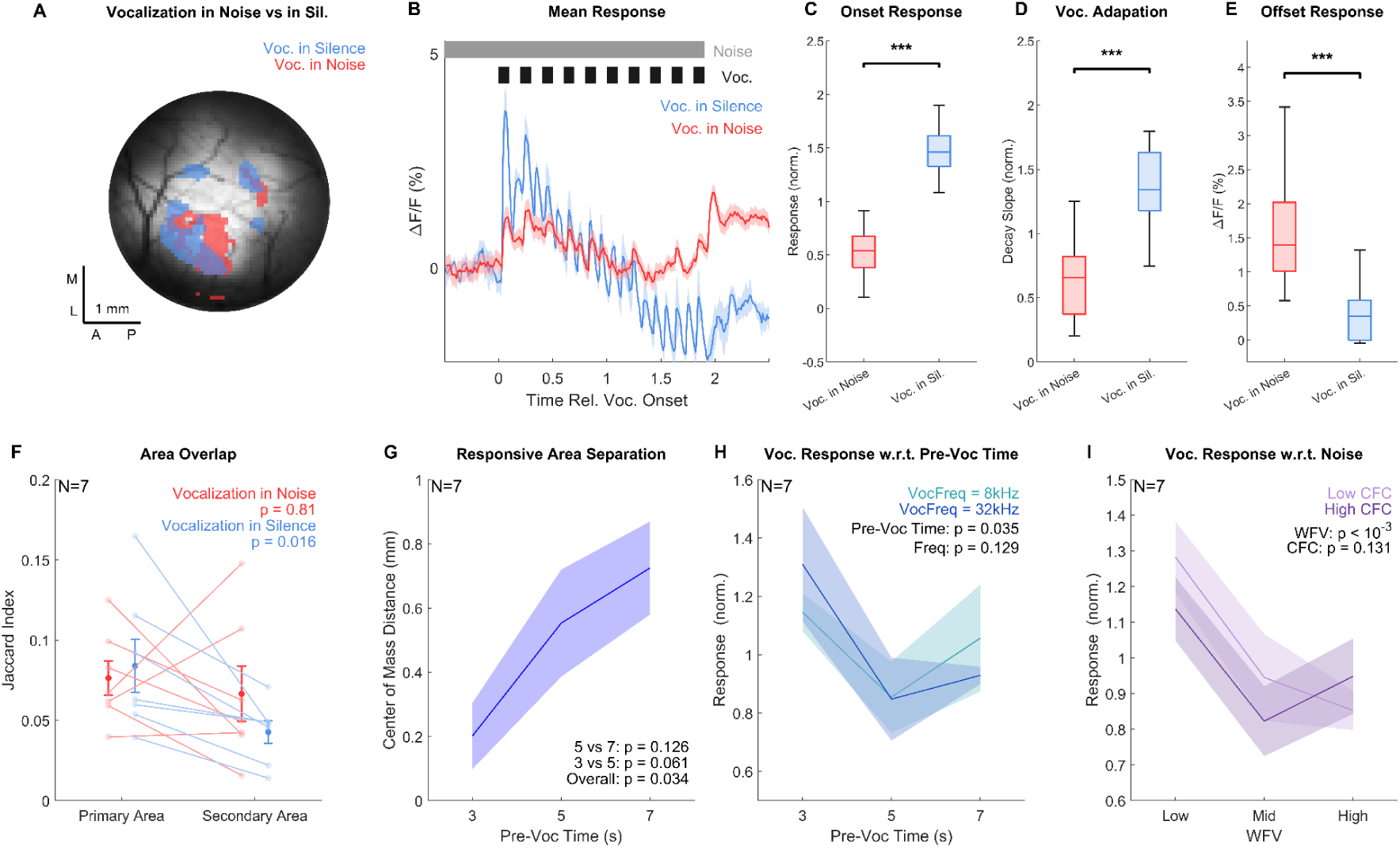
Cortical responses to vocalizations differ between noise and silent contexts. **A** Spatial distribution of vocalization responses during noise and silence in a representative animal. The vocalization-in-noise map (red) and vocalization-in-silence map (blue) were identified from z-score–thresholded (z > 1.5) vocalization response maps. **B** Mean fluorescence responses to vocalizations across all structured noise conditions and silence background trials respectively, extracted from a shared vocalization-responsive region, from the same animal in **A**. Responses are aligned to the vocalization onset to allow comparison across different pre-vocalization times. Gray bar: ongoing noise background; black bars: vocalizations. **C** Vocalization responses were substantially larger in silence than in noise (p < 0.001, WSRT). Onset response was defined as the mean within 400 ms after vocalization onset. For **C**–**E**, the 3-, 5-, and 7-s pre-vocalization time conditions were pooled across animals (7 animals × 3 conditions). **D** Responses to vocalizations adapted significantly more strongly in silence (p < 0.001, WSRT). Decay slopes were estimated from first-order fits to region-averaged responses and used to quantify adaptation strength, with more negative values indicating stronger attenuation. Slopes were normalized by the mean slope across paired conditions within each animal. **E** Offset-response amplitude was quantified as the rise from the local post-offset trough to the subsequent response peak within 300 ms after stimulus termination. The offset response was significantly smaller in silence than in noise (p < 0.001, WSRT). **F** Overlap of vocalization-in-noise (red) and vocalization-in-silence (blue) response maps with primary and secondary auditory fields, quantified using the Jaccard index. Paired primary–secondary comparisons showed significantly greater primary overlap for vocalizations in silence (p = 0.016, WSRT), but not for vocalizations presented during ongoing noise (p = 0.81, WSRT). **G** The spatial separation between the vocalization-in-noise (red) and vocalization-in-silence (blue) responsive maps. Spatial separation was measured as distance between their centers of mass, increased with pre-vocalization noise duration (LME model with fixed effect onset time and random effect animal). **H** Vocalization response magnitude as a function of pre-vocalization time and vocalization frequency (two-factor LME model with fixed effect onset time and vocalization frequency, with animal as a random effect). Response magnitude was quantified as the 90th percentile of the vocalization-evoked activity map and normalized within animal. **I** Vocalization response magnitude as a function of noise statistics (two-factor LME model with WFV and CFC as fixed effects and animal as a random effect).

This similarity was further assessed by quantifying overlap with classically defined auditory areas based on response latency and tonotopy. For vocalizations presented in silence, the response map overlapped significantly more strongly with primary (A1/A2/AAF) than secondary auditory cortex (DP/DM/TeA; p = 0.016, WSRT). This primary bias was absent for vocalizations presented during ongoing noise (p = 0.81, WSRT; Fig. 3F).

In contrast, response dynamics differed markedly. Responses were extracted from a shared vocalization-responsive region (see Methods) to ensure that differences reflected stimulus context rather than spatial selection. Vocalizations in silence and in noise exhibited distinct response profiles (Fig. 3B from an example animal, qualitatively consistent across animals). Compared to noise conditions, vocalizations presented in silence showed stronger onset responses (normalized response 1.5 ± 0.1 vs 0.55 ± 0.1; p < 0.001, WSRT; Fig. 3B, 3C), steeper decay following the peak (p < 0.001, WSRT; Fig. 3D), and a smaller offset response at the end of the stimulus (p < 0.001, WSRT; Fig. 3E). The larger offset response in the noise condition is consistent with the simultaneous termination of the noise background and the final vocalization.

These differences indicate that cortical responses to vocalizations depend strongly on the ongoing acoustic context, affecting both their temporal dynamics and spatial organization.

Although the overall spatial localization was similar, the relative positioning of the response regions diverged with increasing pre-vocalization duration. The distance between the centers of mass of the vocalization-in-noise and vocalization-in-silence response areas increased as the duration of preceding noise increased (p = 0.034, LME; Fig. 3G), suggesting that these differences may be related to adaptation or suppression induced by the preceding noise, leading to changes in the spatial representation of vocalizations.

*Vocalization responses in noise depend on the statistical structure of the preceding stimulus* We next asked how the properties of the preceding noise influence vocalization responses. Focusing on the vocalization-in-noise condition, response magnitude varied systematically with both temporal context and stimulus statistics.

Response strength decreased with the duration of preceding noise (p = 0.035, two-factor LME; Fig. 3H), indicating an effect of stimulus history on vocalization responses. In addition, responses were strongly modulated by within-frequency variability (WFV) (p < 0.001, two-factor LME; Fig. 3I), with larger responses observed for low WFV stimuli. In contrast, cross-frequency correlations (CFC) had a weaker, non-significant effect (Fig. 3I).

These results indicate that vocalization responses in noise are not fixed, but are contextually shaped by the statistical structure of the ongoing sensory environment.

### Decoding of vocalizations improves with preceding noise exposure

Previous research has suggested that increasing exposure duration improves both behavioral discrimination (McDermott et al., 2013) and cortical representation (Alishbayli et al., 2025). Above we find that the response to vocalizations decreased with exposure duration in size (Fig. 3H) but became more separated in space (Fig. 3G). As this provides no clear conclusion as to whether the stimulus representation improves, we turned to a more direct method of assessing this, neural decoding using Support Vector Machines, mirroring our previous approach (Alishbayli et al., 2025). We trained an SVM to classify neural activity during noise-sustain periods, defined as the 1 s period preceding vocalization onset, versus vocalization-onset periods, defined as the 10×100 ms vocalization epochs within the subsequent vocalization sequence. Each sample consisted of the spatial pattern of activity across pixels at a given time point.

We first contrasted decoding from two regions of AC: primary auditory cortex, represented by an ROI centered on A1 and defined independently based on tonotopy and latency, and a vocalization-responsive region, defined from responses to vocalizations embedded in noise (Fig. 4A). For this region-specific analysis, circular ROIs of matched size were centered within the corresponding regions, ensuring comparable SVM input dimensionality. In an example trial, responses from the two ROIs followed broadly similar stimulus-locked temporal dynamics, but differed in response magnitude and pixel-wise response structure (Fig. 4B).

**Figure 4.**
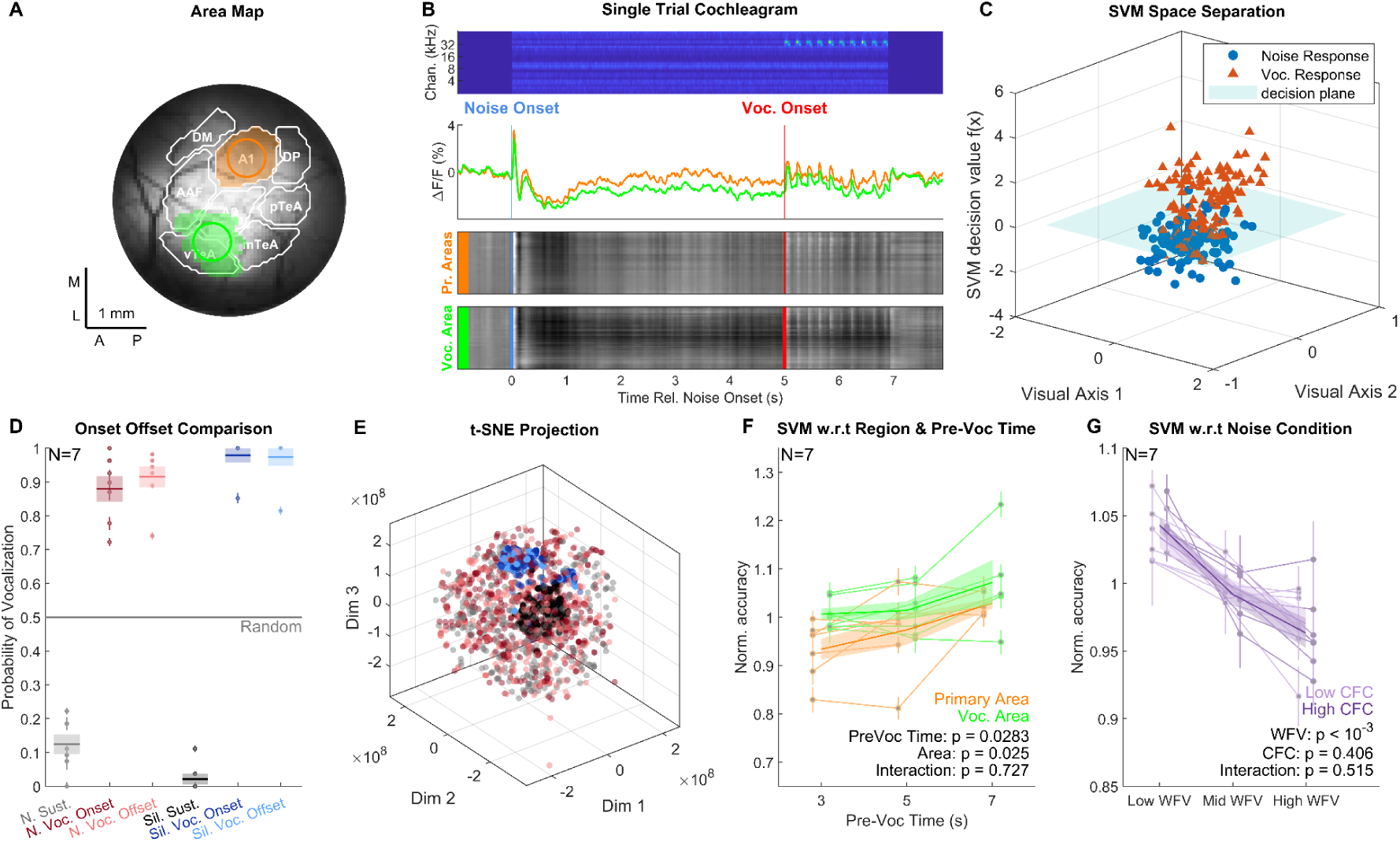
Decoding of the presence of vocalizations in the context of structured noise. **A** Spatial locations of matched circular ROIs in a representative animal: an A1-centered ROI (orange) and a vocalization-responsive ROI (green). A1 was identified using tonotopy and latency, whereas the vocalization-responsive region was identified from responses to vocalizations embedded in noise. Classical auditory areas are indicated by white contours. **B** Example responses from the two ROIs for a single trial. Top, cochleogram of the stimulus; middle, mean fluorescence responses within each ROI; bottom, pixel-wise responses sorted by response similarity. **C** Representative visualization of the SVM classifier used to distinguish noise-sustain from vocalization-onset activity. Noise-sustain samples were drawn from the 1 s period preceding vocalization onset, and vocalization-onset samples from the 10×100 ms vocalization periods within the subsequent sequence (1 s total). Neural activity is projected into a low-dimensional space for visualization; the plane at Z = 0 indicates the SVM decision boundary, and points are colored according to their true labels. **D** Generalization of decoding across temporal segments of the vocalization sequence using activity from the entire craniotomy. The classifier was tested on pre-vocalization sustain, vocalization-onset, and inter-vocalization offset (pause) periods. Decoder output is expressed as the probability of assignment to the vocalization class, with higher values indicating stronger assignment to the vocalization-onset class. Results are shown separately for vocalization-in-noise (light grey, dark red and light red for each segment respectively) and vocalization-in-silence (black, dark blue and light blue for each segment respectively) trials. Points represent individual animals (N = 7); bars indicate mean ± SEM, and the gray line indicates chance level. **E** t-SNE visualization of response patterns (reduced to 3 dimensions) using activity from the entire craniotomy. Each point represents one frame from a condition-averaged response and is labeled according to stimulus context and temporal segment: pre-vocalization sustain, vocalization-onset, or inter-vocalization offset periods in noise or silence trials. Same corresponding colors were used for each condition as in **D**. **F** Normalized decoding accuracy as a function of pre-vocalization time for matched A1-centered and vocalization-responsive ROIs. Accuracy increased with pre-vocalization time and remained higher in the vocalization-responsive ROI. Statistics were assessed using a linear mixed-effects model with cortical ROI and pre-vocalization time as fixed effects and animal as a random effect (N = 7 mice); shaded error regions indicate ±1 SEM across animals. **G** Normalized decoding accuracy across structured noise conditions using activity from the entire craniotomy. Accuracy is shown as a function of WFV and CFC. Statistics were assessed using a linear mixed-effects model with WFV and CFC as fixed effects and animal as a random effect (N = 7 mice); error regions indicate ±1 SEM across animals.

To illustrate the decoding principle, population activity was projected into a low-dimensional space, in which activity during vocalization-onset periods could be separated from activity during noise-sustain periods by the SVM decision boundary (Fig. 4C). This visualization is shown for demonstration purposes, while classification itself was performed directly on the high-dimensional spatial activity patterns.

We next examined whether the decoder output generalized across temporal segments of the vocalization sequence using activity from the entire craniotomy (Fig. 4D). The classifier was trained to distinguish noise-sustain from vocalization-onset periods and then evaluated on pre-vocalization-sustain, vocalization-onset, and inter-vocalization offset (pause) periods. For intuitive interpretation, we expressed decoder output as the probability of assignment to the vocalization class, with chance level at 0.5 (gray line). This measure was derived from classification accuracy: for vocalization-onset periods it corresponded directly to the fraction classified as vocalization, whereas for noise-only periods the inverse classification accuracy was used, such that higher values consistently indicated stronger assignment to the vocalization class. In both vocalization-in-noise and vocalization-in-silence trials, pre-vocalization sustain periods were predominantly assigned to the non-vocalization class, whereas vocalization-onset periods were strongly assigned to the vocalization class. Notably, activity during the offset (pause) periods between vocalizations was also strongly assigned to the vocalization class, despite the absence of vocalization sound during these periods. This indicates that vocalization-related cortical activity persists beyond the acoustic epochs themselves.

To further explore the structure underlying this temporal persistence, we visualized condition-averaged population responses from the entire craniotomy using three-dimensional t-SNE (Fig. 4E). In silence trials, activity patterns of pre-vocalization periods showed relatively clear separation against post-vocalization periods (both onset and offset) in the low-dimensional embedding. In contrast, the corresponding activity patterns during structured-noise trials were more intermixed in this representation. This suggests that vocalization-related activity in the presence of structured noise is embedded in a more complex population response structure, for which only high-dimensional decoding approaches such as SVMs provide a useful readout.

We then tested whether decoding performance depended on the duration of preceding noise exposure using the matched A1-centered and vocalization-responsive ROIs (Fig. 4F). Normalized decoding accuracy improved with pre-vocalization time, from 0.93 ± 0.03 to 1.04 ± 0.03 in primary area (A1-centered) ROI and from 1.00 ± 0.02 to 1.07 ± 0.03 in the vocalization-responsive ROI between 3 and 7 s, and was consistently higher in the vocalization-responsive ROI. A linear mixed-effects model with pre-vocalization time and cortical ROI as fixed effects and animal as a random effect revealed significant effects of pre-vocalization time (p = 0.028) and ROI (p = 0.025), with no significant interaction between the two factors (p = 0.727; N = 7 mice). Thus, longer exposure to the structured-noise context improved the discriminability of subsequent vocalization-related activity, while the vocalization-responsive region ROI provided more decodable information than A1-centered ROI.

Finally, we evaluated decoding performance across the statistical structure of the background noise using activity from the entire craniotomy (Fig. 4G). Normalized decoding accuracy decreased with increasing within-frequency variance (WFV), from 1.04 ± 0.01 at low to 0.96 ± 0.01 at high WFV. A linear mixed-effects model with WFV and cross-frequency correlation (CFC) as fixed effects and animal as a random effect confirmed a significant effect of WFV (p < 0.001), but revealed no significant effect of CFC (p = 0.406) and no significant WFV-by-CFC interaction (p = 0.515; N = 7 mice). Thus, vocalizations became less discriminable from the ongoing noise as its within-frequency variance increased, whereas CFC had little systematic influence on decoding.

Together, these results show that cortical representations of vocalizations become progressively more discriminable with ongoing exposure to background noise and are modulated by the within-frequency variance of that background, while exhibiting temporal persistence that extends beyond the acoustic signal. The latter appears related to a recent result highlighting that offset responses in the auditory cortex can be informative about stimulus identity (Lamothe et al., 2025).

### Stimulus reconstruction quality depends on both stimulus statistics and time

To assess how much acoustic information is overall preserved in cortical population activity, we trained a linear model to reconstruct the cochleogram representation of structured noise stimuli from widefield responses (Mesgarani et al., 2009; Keine et al., 2017). In this process, the neural activity is transformed into a lagged design matrix and used to predict cochleogram features using linear, time-neuron filters (Fig. 5A).

**Figure 5.**
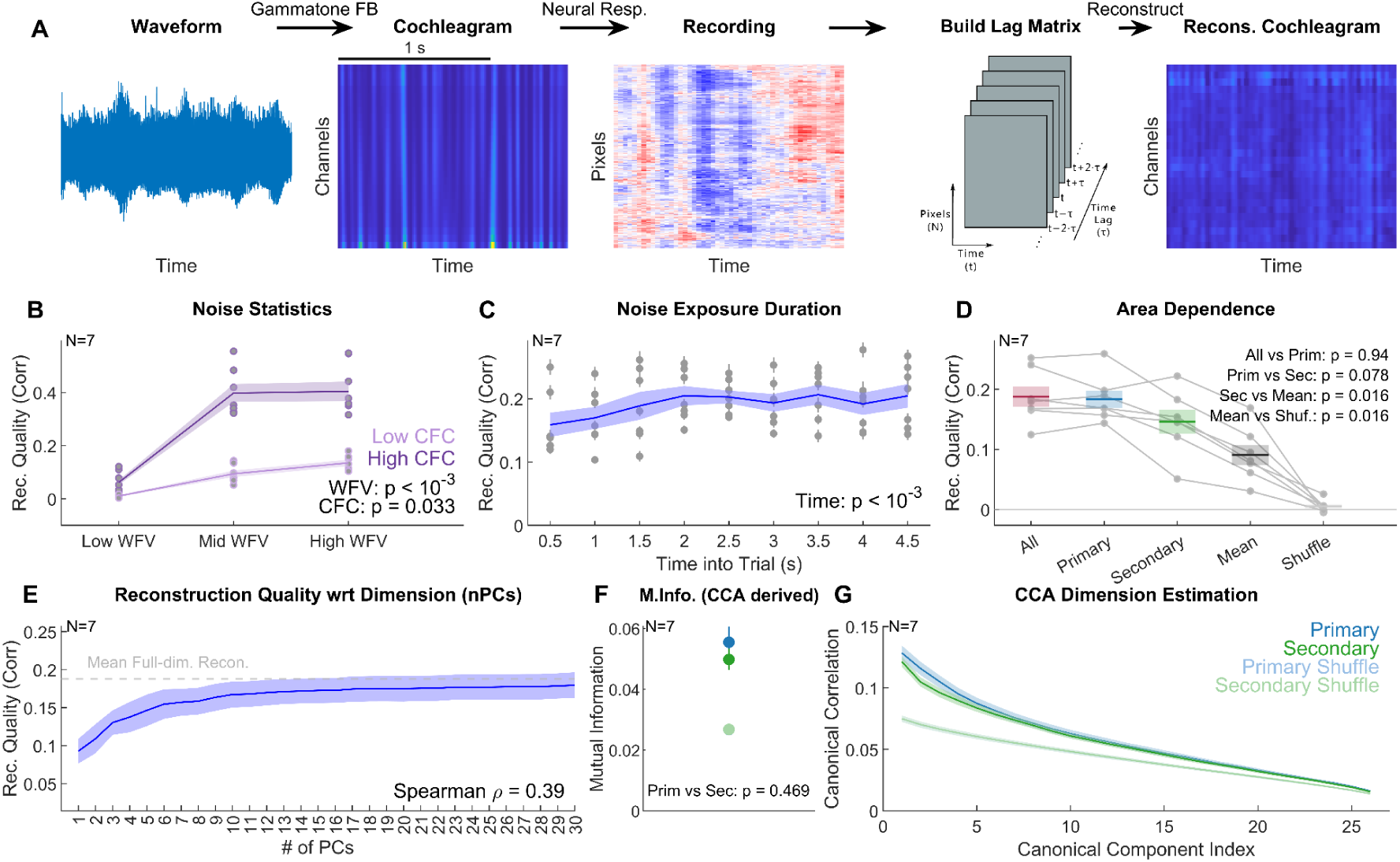
Reconstruction of structured noise from cortical population activity. **A** Reconstruction pipeline. Acoustic waveforms were transformed into cochleograms using a gammatone filterbank. Neural population responses in response to structured noise were organized into a design matrix with lags spanning [−0.2,0.3] seconds relative to each cochleogram time bin. A linear, ridge regression model was trained to predict cochleogram channels from neural activity. **B** Reconstruction quality significantly improved with WFV and CFC (LME model with animal as a random effect, N = 7 animals). Reconstruction quality defined as Pearson correlation between reconstructed and original cochleogram channels, and error hulls denote 1 SEM across animals. **C** Reconstruction quality also improved significantly as a function of time after noise onset, computed within 1-s windows advancing in 0.5-s steps following noise onset (LME model with animal as a random effect, N = 7 animals). Only trials with at least 5 s of noise before the vocalization were included. **D** Area dependence of reconstruction performance. Reconstruction was performed using matched numbers of randomly sampled pixels spanning all auditory areas (red), primary auditory cortex (A1/A2/AAF; blue), or secondary auditory cortex (TeA/DM/DP; green), together with a mean-response control (dark gray) and temporally shuffled control (light gray). Points represent animal-level values and bars indicate mean ± SEM (N = 7). Pairwise statistics are shown in the panel. **E** Reconstruction quality as a function of neural dimensionality. Neural responses were projected onto increasing numbers of principal components before reconstruction. The curve shows mean ± SEM across animals (N = 7). The dashed gray line indicates mean reconstruction performance of the corresponding full-dimensional all-area neural population. Reconstruction increased rapidly over the first several components and approached full-dimensional performance by approximately 10 PCs. **F** CCA-derived information between neural responses and cochleogram features in primary and secondary auditory cortex and their temporally shuffled controls. Identical numbers of pixels were used for both areas and across animals (50 pixels). Points indicate group means ± SEM across animals (N = 7). CCA-derived information was numerically higher in primary cortex, but the primary–secondary difference was not significant (WSRT, p = 0.469). **G** Canonical correlation spectra between neural population activity and cochleogram features for primary and secondary auditory cortex and temporally shuffled controls. Curves indicate mean ± SEM across animals (N = 7). Canonical correlations were strongest in lower-order components and decreased with component index; primary cortex showed slightly higher values than secondary cortex among several lower-order components.

Reconstructed cochleograms moderately resembled the original stimulus representation, with the highest average correlation reaching ∼0.4 among all structured noise conditions, demonstrating that cortical population activity captured via widefield imaging and fast Ca-indicators contains sufficient information to recover spectrotemporal features of the sound.

Reconstruction accuracy varied systematically with both stimulus statistics and the duration of ongoing stimulation. Higher WFV and CFC led to improved reconstruction performance (Fig. 5B; two-factor LME, WFV: p < 0.001, CFC: p = 0.033). Under high CFC, reconstruction correlation increased from 0.06 ± 0.02 at low to 0.40 ± 0.04 at medium WFV, but changed little from medium to high WFV (0.41 ± 0.04), whereas under low CFC it increased more gradually (0.01 ± 0.01, 0.10 ± 0.02 and 0.14 ± 0.02 for low, medium and high WFV respectively), suggesting diminishing returns at higher stimulus structure (see Discussion for a critical interpretation of these results). Reconstruction accuracy also mildly increased over time following noise onset (Fig. 5C; LME, p < 0.001), with improvements mainly occurring at the beginning around the onset dynamics. At later time points beyond ∼2 s, the performance appears to saturate (see Discussion).

### Reconstruction relies on distributed and spatially structured population activity

We next examined how reconstruction performance depended on the spatial extent and cortical location of the neural activity used for reconstruction (Fig. 5D). To control for population dimensionality, reconstruction from all auditory areas, primary auditory cortex, and secondary auditory cortex was performed using matched numbers of randomly sampled pixels within each animal. Reconstruction using pixels sampled across the auditory cortex was comparable to reconstruction from primary auditory cortex (WSRT, p = 0.94), whereas reconstruction was higher in primary than secondary auditory cortex in 6 of 7 animals, showing a tendency toward higher reconstruction performance in primary cortex (WSRT, p = 0.078). Nevertheless, reconstruction from secondary auditory cortex remained substantially above the mean-response control (WSRT, p = 0.016), and the mean-response control exceeded temporally shuffled data (WSRT, p = 0.016). In a control analysis, allowing the all-area condition to use a larger number of pixels did not show a clear trend of improved reconstruction performance, indicating that the comparable all-area and primary performance was not simply imposed by matching population dimensionality. These results indicate that acoustic information is broadly distributed across auditory cortex, with a modest tendency towards higher reconstruction performance in primary than secondary auditory cortex.

*Neural–stimulus relationships are distributed across dimensions and support reconstruction* To test how neural dimensionality contributes to reconstruction, we projected population responses onto progressively larger numbers of principal components (PCs) before fitting the reconstruction model. Reconstruction quality increased rapidly over the first several components and approached the performance of the full-dimensional representation by approximately 10 PCs (Fig. 5E). Additional components produced only modest improvements. Although reconstruction continued to increase overall with dimensionality (Spearman’s ρ = 0.39), most reconstruction-relevant information was therefore captured by a low-dimensional subspace (∼10 PCs) relative to the full pixel-wise representation.

To further characterize the shared structure between neural activity and the acoustic stimulus, we performed canonical correlation analysis (CCA) between population responses and cochleogram representations. CCA identifies successively weaker pairs of neural and stimulus feature combinations whose temporal activity is maximally correlated. Canonical correlations were strongest for the first few components and decreased progressively with increasing component index (Fig. 5G). The primary auditory cortex showed slightly higher canonical correlations than secondary auditory cortex among the lower-order components, although the difference between the two regions was small.

Consistent with this descriptive difference, the mean CCA-derived information estimate was numerically higher in primary than secondary auditory cortex (Fig. 5F). However, this difference was not statistically significant across animals (WSRT, p = 0.469), indicating that the present analysis does not provide evidence for a robust regional difference in total CCA-derived information. Notably, unlike the lagged reconstruction analysis, CCA captures shared neural–stimulus structure at corresponding time points without explicitly incorporating temporal context. The weak regional difference observed with CCA is therefore not necessarily inconsistent with the stronger primary–secondary difference observed for reconstruction, which can additionally exploit temporally distributed stimulus information. Both regions nevertheless showed clear neural–stimulus correspondence relative to shuffled controls.

These analyses indicate that acoustic information is spatially distributed across auditory cortex while much of the information relevant for linear reconstruction is concentrated within a relatively small number of population dimensions. Primary cortex showed a modest numerical advantage in several measures, but shared stimulus-related information was prominent in both primary and secondary areas.

## Discussion

In this study, we used fast widefield calcium imaging to examine how mouse auditory cortex represents structured acoustic backgrounds and vocalizations embedded within them. Structured noise evoked rich cortical dynamics that depended on its spectrotemporal statistics, with early onset responses, later suppressive components, and ongoing fluctuations showing distinct spatial biases across auditory cortical fields. Reconstruction analyses further showed that information about the background was broadly distributed across auditory cortex and became more reliable after the initial onset and early adaptation period, before reaching a more stable encoding state. Against this evolving background representation, vocalizations presented in noise and silence activated partially overlapping cortical regions but differed in response dynamics, spatial organization, and population discriminability. In particular, vocalization response magnitude did not increase but showed a decreasing trend with longer exposure to the preceding noise, whereas decoding of vocalization-related activity improved with exposure time. This dissociation suggests that recent acoustic context can reshape the population representation of embedded vocalizations without necessarily increasing the average evoked response. Together, these findings indicate that the auditory cortex represents both background statistics and target sounds through distributed, context-dependent activity that evolves over time.

### Background context reshapes vocalization representations beyond response magnitude

A central implication of these results is that the influence of background noise on vocalization responses cannot be described simply as masking or attenuation. Vocalizations presented in structured noise produced responses that differed from responses to the same vocalizations in silence, both in temporal profile and in spatial organization. In silence, responses following individual vocalizations also differed in shape from the response after the final vocalization. However, the effect of preceding noise exposure was not captured by response magnitude alone. Vocalization response strength did not show an increase with longer exposure to the background, whereas population decoding improved with pre-vocalization exposure time. This dissociation suggests that recent acoustic context may change the structure of population activity in a way that makes vocalization-related patterns more separable, even when the average evoked response is not enhanced. In this sense, adaptation to background statistics may not only suppress ongoing or redundant components of the stimulus, but may also reshape the cortical state in which an embedded target is represented. Consistent with this, reconstruction of the background did not decline with longer exposure, as would be expected from simple suppression, but improved over the first seconds and then remained stable (Fig. 5C). This interpretation is in line with behavioral and electrophysiological work showing that longer exposure to structured backgrounds can improve vocalization detection and neural encoding (Alishbayli et al., 2025), as well as with human studies showing that auditory cortex can adapt to background noise while improving the representation of foreground speech (Khalighinejad et al., 2019). Our results extend this idea by showing that such context-dependent changes are visible in distributed cortical population activity even during passive listening.

### Spatial response biases coexist with distributed stimulus information

The relationship between response maps and reconstruction performance further suggests that auditory cortical representations are spatially biased but not strictly localized. Previous work has described auditory cortex as combining large-scale field organization with substantial local heterogeneity, and has shown that primary, higher-order, and association regions can differ in their response properties and in the format of complex-sound representations (Feigin et al., 2021; Kanold et al., 2014; Romero et al., 2020). In line with this view, we found that structured-noise onset responses were biased toward primary auditory fields, whereas later suppressive components did not show a significant primary–secondary bias. Reconstruction analyses nevertheless showed that acoustic information was not confined to a single cortical subdivision. When the number of sampled pixels was matched across regions, reconstruction using activity sampled across auditory cortex was comparable to reconstruction from primary auditory cortex, whereas reconstruction from secondary cortex tended to be lower. Allowing the all-area condition to use a larger number of pixels did not significantly improve reconstruction performance, suggesting that the similarity between all-area and primary reconstruction was not simply imposed by matching population dimensionality.

CCA provided a complementary view of this regional organization. Primary cortex showed slightly higher canonical correlations than secondary cortex, particularly among lower-order components, and a numerically higher CCA-derived information estimate, although the latter difference was not significant across animals. This need not mirror the reconstruction result directly, because the two analyses emphasize different aspects of the neural–stimulus relationship: reconstruction explicitly incorporates neural activity across temporal lags, whereas the present CCA measures contemporaneous covariance between neural population activity and cochleogram features. Stimulus information expressed with different delays or temporal dynamics across cortical regions could therefore contribute similarly to lagged reconstruction while producing different instantaneous neural–stimulus correlations. Together, these findings suggest that primary and secondary auditory cortex both contain substantial acoustic information, while differing modestly in the strength and temporal organization with which that information is expressed.

*Within-frequency variance and cross-frequency correlation shape different cortical readouts* The effects of within-frequency variance and cross-frequency correlation indicate that different forms of acoustic structure shape cortical activity in different ways. Both features are related to sound texture statistics, which have been proposed to capture perceptually relevant structure in naturalistic sounds and to influence neural responses in the auditory midbrain and cortex (McDermott et al., 2013; McDermott & Simoncelli, 2011; Mishra et al., 2021; Peng et al., 2024). In our data, WFV had broad effects on cortical dynamics, including ongoing response fluctuations, recovery from suppression, vocalization response magnitude, and decoding performance. This is consistent with studies showing that auditory cortical responses adapt to recent contrast or fluctuation statistics (Cooke et al., 2018; Rabinowitz et al., 2011). CFC, by contrast, contributed more clearly to local response coherence and stimulus reconstruction, but had weaker or less consistent effects on vocalization decoding. The reconstruction result should, however, be interpreted cautiously, as stimulus statistics can influence not only the neural representation but also the intrinsic predictability of the stimulus being reconstructed. In particular, stronger cross-frequency correlations impose additional structure and redundancy across channels, which may make the cochleogram easier to predict and could therefore partially contribute to the improved reconstruction performance.

One possibility is that CFC mainly shapes the representation of the ongoing background by coordinating activity across frequency channels and cortical regions, whereas WFV more directly determines the magnitude of within-channel fluctuations and the adaptation state into which the vocalization is embedded. The weaker CFC effect on vocalization decoding therefore does not imply that CFC is unimportant, but suggests that its influence may be more apparent in measures of background structure than in passive decoding of vocalization presence. This interpretation is partly different from a related behavioral and electrophysiological study, where high CFC slightly, but significantly improved vocalization detection and neural encoding (Alishbayli et al., 2025), but the difference does not necessarily imply a contradiction. CFC-dependent effects may depend on behavioral engagement, the temporal precision and locality of electrophysiological recordings, or sampling from specific auditory fields, whereas the present widefield analysis emphasizes broader population-level activity during passive listening.

### Limitations and future improvements

The present stimulus-design was motivated by the intent to stay close to naturally occurring textural sounds while maintaining precise control over the spectral and temporal properties of the sounds. This approach differed from our previous study (Alishbayli et al., 2025), where naturally occurring sound statistics were combined to create naturalistic sounds. Neither of these approaches in generating sounds leads to stimuli that sound truly natural, as was the original intent of the texture synthesis toolbox (McDermott & Simoncelli, 2011). Full stimulus control and fully natural stimuli obviously cannot be achieved simultaneously. Whether natural sound textures are processed differently because of prior familiarity therefore remains an open question. This could be tested by exposing mice in their home cage to behaviorally relevant sound textures and comparing their responses with those of unexposed animals.

Secondly, it would be interesting to study the presently detected differences also on the cellular level using 2-photon imaging in the same animals. This would provide the possibility to overcome the local averaging that is inherent to widefield imaging (Romero et al., 2020; Waters, 2020). This could highlight subpopulations of cells that show different types of temporal dynamics and potentially integration.

Lastly, the influence of behavioral state and attention could be examined by repeating the present experiments in a task in which mice actively detect vocalizations for reward. This would allow comparison with passive listening and with previous behavioral work (Alishbayli et al., 2025).

## Materials & Methods

### Experimental Model and Subject Details

All experiments were performed at the Central Animal Laboratory of the Radboud University Medical Center. Experimental procedures and protocols were approved by the Dutch Central Commission for Animal Research (*Centrale Commissie Dierproeven*; project numbers 2017-0041 and 2023-0031) and were conducted in accordance with the local animal welfare body. Adult female CBA/JRj mice (Janvier & Charles River Labs) were group-housed under a standard 12 h light/dark cycle with ad libitum access to food and water. This strain was used because CBA/JRj mice retain normal hearing over several months, in contrast to C57BL/6-derived strains, which develop early age-related high-frequency hearing loss (Romero et al., 2020; Zheng et al., 1999). Surgical procedures were performed when animals were 6–8 weeks old, and all recordings included in this study were obtained within 3 weeks after surgery.

### Implant and Transfection Surgery

Mice received pre-operative analgesia via the drinking water in the 24 h preceding surgery (Carprofen, 5 mg/kg, estimated based on regular water intake). At the start of surgery, animals were anesthetized with isoflurane in a 1:1 oxygen/air mixture using 3% isoflurane for induction and 1–2% for maintenance. Respiratory rate was monitored throughout the procedure and maintained between 40 and 80 breaths per minute. After induction of anesthesia, Carprofen (5 mg/kg) and Dexamethasone (2 mg/kg) were administered i.p. Core body temperature was maintained using a homeothermic blanket (RWD RS-485) with feedback from a rectal thermometer, and sterile saline was administered i.p. at 0.5 ml/h.

After induction and preparation for surgery, the scalp was cleared of fur with depilatory cream and disinfected sequentially with Betadine and 70% ethanol. A small volume of local anesthetic, consisting of a Lidocaine/Bupivacaine mixture (0.1 ml), was injected subcutaneously using a fine syringe needle. The skull was accessed through a midline incision. On the right side, the periosteum was removed with a spudger and 3% H₂O₂. The exposed skull surface was lightly roughened with a scalpel to promote adhesion of the dental cement. A custom aluminum headpost was then attached to the right skull hemisphere with dental cement (Superbond, Sun Medical), positioned at approximately 45° relative to the medial axis. After headpost implantation, the left side of the skull was exposed down to the ventral temporal bone and cleaned with 3% H₂O₂.

The craniotomy was targeted to the auditory cortex in the left hemisphere. Its estimated center was placed 2.4-3 mm posterior and 4.7–5.0 mm lateral to Bregma, approximately 0.5 mm medial to the parietal-temporal ridge. These coordinates were measured relative to Bregma using a 650 nm, 5 mW crosshair laser (Quiaoba) mounted perpendicular to the skull surface on a micropositioner (Kopf Instruments). Before opening the skull, the surrounding bone was cleaned, roughened, and reinforced with dental cement. Depending on the final window size, a circular craniotomy of 3–4.3 mm diameter was made around the estimated center. Bone removal was performed using a micromotor drill (K.1070-2, Foredom) fitted with a 0.5 mm steel burr (19007-05, Fine Science Tools), together with #4 forceps (Fine Science Tools). To reduce the risk of damaging the dura or surface vasculature, a small handle was first cemented to the central bone piece, which allowed the central bone disc to be lifted out gently. Once the craniotomy was opened, the exposed dura was kept hydrated by repeated application of sterile saline.

For viral expression of the calcium indicator jGCaMP8m, injections were performed with a pulled glass micropipette made from a 5 μl capillary, with an unbevelled inner tip diameter of approximately 20 μm (555/5, Assistent). The pipette was back-filled with mineral oil and loaded with approximately 2.5 μl of viral solution containing pGP-AAV9-syn-jGCaMP8m-WPRE (#162375, Addgene; 2 × 10¹² vg/ml). Using a Sensapex SMX micromanipulator, the pipette was advanced at an angle of approximately 45° relative to the brain surface. Injection coordinates were shifted medially such that the final injection pattern formed an 800–1000 μm-spaced triangular tiling centered on the craniotomy. Viral solution was delivered at approximately 400 μm cortical depth, corresponding to a 560 μm travel distance along the angled pipette trajectory. Each injection site received 150–250 nl of virus at a rate of approximately 30–60 nl/min. After each injection, the pipette was left in place for 4 min before withdrawal to limit backflow.

The cranial window consisted of two smaller circular glass coverslips, 3, 3.5, or 4.3 mm in diameter, bonded to one larger coverslip, 4, 4, or 5 mm in diameter, using optical adhesive (NOA68, Norland; #0 thickness glass, Warner Instruments). Before implantation, the assembled window was sterilized in 70% ethanol and rinsed with double-deionized water. The window was then lowered into the craniotomy and cemented in place while applying gentle pressure to the underlying tissue. To facilitate reliable repositioning across imaging sessions, a custom grade 5 titanium headplate was finally cemented around the preparation. The headplate was aligned with the skull midline using the micropositioner and positioned planar to the cranial window.

Following surgery, mice were given one week to recover. Habituation to head fixation and the imaging setup began thereafter and continued for at least one additional week, during which sugar water was used as a reward. Starting in the second week after surgery, viral expression was monitored regularly, and experiments were initiated once fluorescence levels had stabilized.

### Widefield Imaging

Widefield imaging was performed in a dark, sound-insulated recording chamber. The inner walls of the chamber were lined with acoustic foam with a black surface coating (50 mm, Basotect Plan50, BASF). For frequencies above approximately 1 kHz, this material provides a sound absorption coefficient greater than 0.95, defined as the fraction of incident sound intensity that is absorbed. This corresponds to more than 26 dB of additional sound attenuation, on top of the acoustic shielding provided by the chamber itself.

Fluorescence excitation was provided by a 470 nm LED (M470L4, Thorlabs), driven by a multi-channel LED driver (DC4104, Thorlabs). The LED was pulsed at 100 Hz with 1 ms pulses at 100 mA. Excitation light was collimated with a lens (ACL2520U, Thorlabs) and focused to approximately 300 µm below the vasculature level through a 4x objective (Plan Fluor 4×/0.13, Nikon), using an illumination intensity of ∼0.38 mW/mm².

Emission light was separated from excitation light using a long-pass dichroic mirror (DMLP490R, Thorlabs) and further filtered with a green band-pass filter (MF525-39, Thorlabs). Images were acquired with a high-speed global-shutter CMOS camera (U3-3060CP-M-GL, IDS Imaging) at 100 frames per second, with an exposure time of 1 ms. Recordings were acquired in a square FOV centered on the craniotomy at a resolution of 1200×1200 pixels and 4×4 spatially averaged inside the camera during acquisition.

To compensate for animal-specific differences in fluorescence intensity, the digital camera gain was adjusted between 1 and 3 such that the maximum raw pixel intensity reached approximately 25–50% of the sensor bit range. Photobleaching was slow under these imaging conditions, remaining below 5% of the total fluorescence intensity over an entire 2–3 h recording session. It was therefore considered negligible over the shorter duration of individual paradigms. Exposure times were acquired on a multichannel data acquisition device (PCIe-6351, National Instruments), sampled at 10 kHz.

### Structured Noise Generation

#### Initial spectrotemporal envelope and waveform generation

Stimuli were constructed in the spectrotemporal domain using a framework based on temporally orthogonal ripple combinations (TORCs), in which envelopes are defined over log-frequency *x* (in octaves) and time t. Each stimulus consisted of a sum of spectrotemporal ripple components:

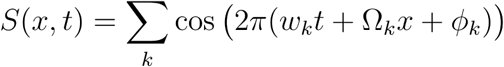

where *w_k_* denotes temporal modulation rate (Hz), Ω*_k_* denotes spectral modulation rate (cycles/octave), and *ϕ_k_* is a random phase. In the implementation, each ripple was generated in a strictly positive form,

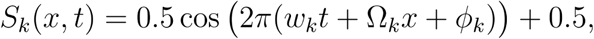

and summed across components to produce the final envelope. Temporal modulation rates were sampled up to 20 Hz with 11 steps. Spectral modulation rates were sampled using 11 steps up to a maximum of 2 cycles/octave for the low-correlation condition and 0.05 cycles/octave for the high-correlation condition. Ripple parameters were arranged on a structured grid with small random perturbations to avoid exact periodicity and to ensure approximate orthogonality across components (Englitz et al., 2010).

The envelope was defined over a log-frequency axis spanning *X*=5 octaves with resolution Δ*x*=0.2 octaves (yielding 26 channels), and over time with a sampling rate of 250 kHz. After summation, the envelope was shifted to be strictly non-negative and scaled such that its mean amplitude was normalized:

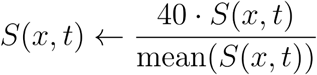

The choice of Δx=0.2 ensures sufficient resolution to represent the highest spectral modulation rates without aliasing, analogous to a Nyquist constraint in the spectral domain. The spectrotemporal envelope was then converted into a time-domain waveform by modulating a bank of sinusoidal carriers. Carrier frequencies were logarithmically spaced according to

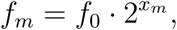

with base frequency *f_0_*=2000 Hz. The final waveform was synthesized as

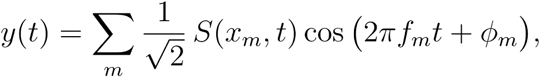

where *ϕ_m_* are random carrier phases. This construction ensures that the imposed spectrotemporal structure directly determines the amplitude envelopes within each frequency channel.

### Measurement of auditory statistics

Stimulus statistics were measured using a model of auditory processing based on cochlear filtering and modulation analysis, following the framework of McDermott and Simoncelli (2011), as implemented in the provided code of their sound synthesis toolbox (https://mcdermottlab.mit.edu/downloads.html).

The waveform *y(t)* was first decomposed using a bank of 36 cochlear-like bandpass filters spanning approximately 500 Hz to 64 kHz. In the TORC configuration, constant-Q cosine filters were used to minimize overlap between adjacent channels. The output of each filter defines a subband signal:

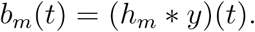

The analytic envelope of each subband was then computed using the Hilbert transform:

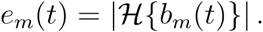

Envelopes were subsequently downsampled from the audio sampling rate (250 kHz) to an envelope sampling rate of 25 kHz to reduce computational cost while preserving modulation structure. Each subband envelope was further analyzed using a modulation filterbank consisting of 100 filters spanning 1–40 Hz. These filters were constructed with approximately constant-Q bandwidth (Q ≈ 10), providing a decomposition of temporal fluctuations at behaviorally relevant modulation rates. For each subband *m* and modulation channel *q*, modulation responses were computed as

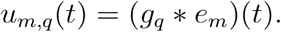

From these representations, several statistics were extracted. For each subband, the envelope mean, standard deviation, and coefficient of variation were computed as:

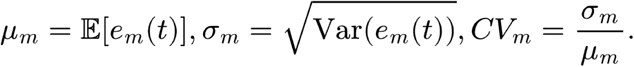

Cross-channel envelope correlations were computed as Pearson correlations between subband envelopes, capturing dependencies across frequency channels. Additional higher-order statistics (e.g., skewness, kurtosis, autocorrelation) were also computed but were not explicitly controlled in the present study.

### Variance adjustment and preservation of correlation structure

Spectral correlation structure was determined during stimulus construction by the choice of spectral modulation range. Specifically, stimuli were generated using either a high spectral modulation condition (2 cycles/octave) or a low spectral modulation condition (0.05 cycles/octave), corresponding to weak and strong correlations across frequency channels. This empirically yields correlations near 0 and ∼0.8 respectively.

To manipulate envelope variability, a nonlinear transformation was applied to the spectrotemporal representation prior to final synthesis. For each subband *m*, the measured envelope variance *σ_m_^2^* was compared to a desired target envelope variance (0.02, 0.2, or 0.4), and an exponent was defined:

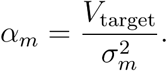

The spectrotemporal envelope in that channel was then transformed according to:

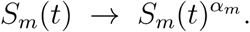

This transformation was applied independently across frequency channels. The modified spectrotemporal representation was subsequently converted back into a time-domain waveform using the same carrier synthesis procedure described above.

Because the transformation is monotonic and applied independently within each channel, it primarily alters marginal statistics (mean and variance) while preserving the temporal structure of fluctuations. As a result, the cross-channel correlation structure imposed by the spectrotemporal ripple components is largely preserved.

### Vocalization waveform construction

The individual vocalization waveform used in the vocalization sequence described above was modelled after a chevron/inverted-u type (Oliveira-Stahl et al., 2023) and generated synthetically as a frequency-modulated sinusoid. Each call had a duration of *L_stim_* =0.1 s and was generated around one of the two base frequencies described below (next section). A discrete time vector was first defined over the call duration,

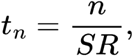

where *SR* is the sound sampling rate and *n* indexes individual sound samples. The instantaneous frequency at each sample was defined as a nonlinear sinusoidal modulation around the base frequency,

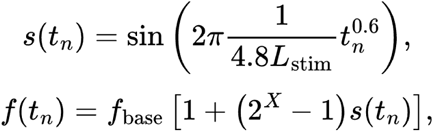

where *f*_base_ is the base frequency and *X*=0.2 octave defines the frequency excursion. The constant 4.8 was used as a fixed shape parameter controlling the curvature and duration of the frequency modulation within the 0.1 s call.

Because the waveform is determined by phase rather than frequency directly, the instantaneous frequency trajectory was converted into a cumulative phase by summing the frequency increment at each sample,

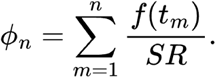

The vocalization waveform was then generated as

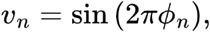

and normalized to unit root-mean-square amplitude. Thus, each call was a 100 ms locally sinusoidal waveform whose instantaneous frequency followed a curved, vocalization-like trajectory around the selected base frequency.

For vocalizations embedded in background sounds, the waveform amplitude was scaled relative to the local background energy around the corresponding base frequency. Specifically, the background was filtered between *f*_base_ and *f*_base_2^X^, and the root-mean-square amplitude of this filtered background was used as the local reference level. The normalized vocalization waveform was then multiplied by this local reference amplitude and by the linear signal-to-noise ratio factor 10^SNR/20^, with SNR= 5 dB. The same waveform-generation procedure was used for vocalizations presented in silence, with amplitude scaling matched to the corresponding frequency-specific reference level.

### Noise-Vocalization paradigm

Each trial began with a structured noise only period lasting 0.5, 1, 3, 5, or 7 s. The noise parameters were varied across conditions, including within-frequency variance (WFV) levels of 0.02, 0.2, or 0.4 and cross-frequency correlation values of 0 or 0.8.

Following the pre-vocalization time, a vocalization sequence was presented on top of the ongoing noise. Vocalizations consisted of 10 repetitions of the vocalization separated by 100 ms silent intervals, resulting in a total vocalization sequence duration of 1.9 s. Vocalizations were generated with base frequencies of either 8 kHz or 32 kHz.

In addition to noise-plus-vocalization conditions, control stimuli consisting of vocalizations presented in silence were also included.

### Noise burst and pure-tone tuning paradigms

Noise-burst and pure-tone tuning paradigms were used as basic reference stimuli for region definition and tuning estimates (see in the following sections). In the noise-burst paradigm, each trial consisted of a 100-ms broadband noise burst presented at 50 dB SPL, followed by a 900-ms pause. Between trials, an additional intertrial interval was randomly varied between 4 and 5 s to reduce temporal predictability. This paradigm provided a basic reference for fast auditory responsiveness across the cortical window.

In the tuning paradigm, 100 ms pure tones were presented at 11 logarithmically spaced frequencies between 2 and 64 kHz and at four sound levels linearly spaced between 20 and 80 dB SPL. Pure tones were presented with a stimulus onset asynchrony (SOA) of 1 s. Each frequency–amplitude combination was repeated six times within a randomized sequence, and each randomization was repeated five times, resulting in 30 presentations per frequency–amplitude combination. Additional recordings using SOAs of 500 ms produced qualitatively similar tonotopic estimates.

## Data analysis

*Widefield image processing:* Frame times were synchronized to the stimulus timing using the camera’s exposure triggers recorded by the data acquisition device. In the rare cases (<0.1%) where a frame was lost during transfer, the missing frame was replaced by the preceding frame, thereby avoiding interpolation from future time points. Images were downsampled by a factor of 2 using MATLAB’s *imresize* function before further processing, resulting in a final image size of 150×150 pixels. All analyses were restricted to pixels within the cranial window.

For each trial, fluorescence signals were converted to ΔF/F₀ on a pixel-by-pixel basis.

The baseline fluorescence, F₀, was defined as the mean fluorescence during the pretrial baseline period for each pixel. This static pretrial baseline was then used to calculate the relative fluorescence change over time. Across recordings, baseline fluorescence could vary between animals and trials, with slow fluctuations observed in both stimulus-driven and silent periods. Over the full recording duration, approximately 1200 s, we observed only a small systematic decrease in baseline fluorescence, which was attributed to a gradual reduction in indicator brightness, possibly due to photobleaching, which, however, did not persist over days.

No spatial or temporal deconvolution was applied, in order to avoid introducing assumptions about spatial light mixing or calcium indicator dynamics into the analysis of spatiotemporal activity patterns. No temporal low-pass filtering was performed, because the onset dynamics of jGCaMP8m contained informative signals up to the full camera sampling rate of 100 fps. To verify temporal alignment and trial sequence reconstruction, simulated video data based on the raw stimulus waveforms were generated for structured noise and vocalization paradigms and processed using the same analysis pipeline. Finally, to reduce the influence of small motion artefacts, typically <1 pixel, on ΔF/F-derived functional measures near vessel boundaries, sub-pixel rigid motion correction was applied using the NoRMCorre package (Pnevmatikakis & Giovannucci, 2017).

### Tonotopy estimate

To improve the estimate of pure-tone-evoked activity, each pixel’s ΔF/F₀ trace was additionally baseline-corrected separately for each frequency–amplitude combination and trial. The baseline was defined as the mean activity during the 200 ms period preceding stimulus onset and was subtracted from the corresponding response trace. Tone-evoked responses were quantified within a 0–230 ms window after stimulus onset. This window included the 100 ms tone presentation and an additional 130 ms to capture the delayed response dynamics of both neural activity and the calcium indicator, based on rise-to-peak and half-decay constants reported previously (Zhang et al., 2023). For each pixel, the peak ΔF/F₀ value within this window was taken as the response amplitude.

Responses were then averaged across trials for each frequency–amplitude combination. For each pixel, frequency–amplitude combinations were retained only when their mean response exceeded the 55th percentile of that pixel’s response distribution across all tested frequencies and amplitudes. Best frequency was estimated following the approach described in Romero et al. (2020). Briefly, the lowest amplitude that produced an above-threshold response was identified as the response threshold for that pixel. Responses were then summed across this amplitude and the next two higher amplitudes, while excluding amplitudes above 70 dB SPL. Across frequencies, a weighted normalized response profile was computed for each pixel, and the frequency with the strongest weighted response was assigned as the pixel’s best frequency.

The overall responsivity of each pixel was quantified as the 55th percentile of its responses across all tested frequencies and amplitudes. Although tonotopic gradients could be recovered, the resulting maps varied considerably across animals and did not always clearly align with the temporal organization of the auditory cortex and surrounding regions. This variability is consistent with previous reports describing inter-animal differences in auditory cortex position and internal subdivision (Narayanan et al., 2023; Romero et al., 2020).

### Region Definitions

Because there is no widely accepted consensus on the precise criteria defining primary and secondary auditory fields (Calhoun et al., 2023; Kanold et al., 2014; Romero et al., 2020; Sawatari et al., 2011), we used a combined, functional approach to delineate auditory areas. In brief, primary-like auditory regions were identified from the low-latency, high-fidelity response center and further subdivided using tonotopic features, whereas surrounding slower-response regions were assigned to secondary auditory fields, including TeA. This approach incorporated conventional tonotopic organization and low-latency response centers together with temporal response fidelity, since secondary auditory regions are generally characterized by slower response dynamics (Kanold et al., 2014). Area borders were defined manually for each animal using a custom-written graphical user interface, following the functional delineation approach described above and previously (Peters et al., 2026).

The use of fast calcium imaging made it possible to resolve temporal response features that are less accessible in GCaMP6-based widefield studies. We therefore treated some posterior tonotopically organized regions as temporal association cortex rather than primary auditory cortex when they exhibited weak tonotopy, long response latency, slower modulation dynamics, and weak impulse responses. This assignment is consistent with classical anatomical labeling in the Allen Brain Atlas and with recent work describing TeA as exhibiting sparser and high-latency auditory responses (Feigin et al., 2021; Wang et al., 2020).

Temporal response fidelity was estimated from responses to single noise burst presentations. For each pixel, we measured the peak-to-trough response distance in ΔF/F₀ during presentation of a 100 ms noise burst at 50 dB SPL. In addition, response timing was quantified by estimating the half-maximum latency around the response peak after Fourier-transform-based interpolation of the trace using MATLAB’s *interpft* function. Together, these measures allowed reliable localization of a set of fast-responding core auditory regions, which we think comprise A1, A2, and AAF. These were distinguished by their short response latencies and rapid rise-to-peak dynamics within the narrow stimulus-evoked time window.

### Identification of structured noise onset and minimum response maps

For the structured-noise response maps, activity was expressed relative to the mean ΔF/F₀ during the pre-noise period (1 s before noise onset). Trials belonging to the same stimulus condition were then averaged to obtain a condition-mean response movie for subsequent computations.

To identify the structured noise onset map, activity was analyzed within an early post-onset window (0-70 ms after noise onset) following noise onset. For each pixel, the maximum ΔF/F₀ value within this window was computed, yielding a map of peak onset responses across auditory cortex.

To identify the structured noise minimum map, activity was analyzed during a later adaptation period (200 ms - 2000ms after noise onset), using only trials with pre-vocalization times ≥ 3 s so that this window did not overlap with vocalizations. For each pixel, the 5th percentile of ΔF/F values within this window was computed, and the minimum response amplitude was defined as the mean of all time points falling below this percentile, providing a robust estimate of the strongest suppression. Response maps were computed separately for each animal and stimulus condition.

Binary noise onset and noise minimum maps (Fig. 2A–F) were obtained by combining the corresponding response maps across pre-vocalization times (mean across all pre-vocalization times for the onset map; median across pre-vocalization times ≥ 3 s for the minimum map), followed by z-score thresholding (z > 1.5).

### Adaptation rate of structured noise responses

Adaptation was quantified on a pixel-by-pixel basis from baseline-subtracted activity. For each pixel, baseline activity was defined as the mean response during the pre-noise period. The peak response amplitude was identified as the maximum ΔF/F₀ value within an early post-onset window (0-70 ms after onset) following noise onset. The subsequent minimum response amplitude was estimated during a later adaptation period (200 - 2000 ms after onset) by computing the 5th percentile of the response values and defining the minimum response as the mean of all time points falling below this threshold. The corresponding minimum time was estimated as the mean time index of these time points. Adaptation slope was then computed for each pixel as the difference between peak and minimum response amplitudes divided by the time elapsed between the peak response and estimated minimum response. Pixels with non-positive or undefined time differences were excluded. Condition-level adaptation values were obtained by averaging pixel-wise adaptation slopes across the cortical mask. For visualization of adaptation-related regions, the sign of the slope map was inverted such that larger positive values indicate stronger adaptation.

### Recovery time of structured noise responses

Recovery was quantified from the mean fluorescence trace extracted from the noise onset map (NOM). For each trace, a 30-ms moving-average filter was applied for signal smoothing prior to feature detection. The trough was defined as the minimum of the smoothed trace within a fixed search window from 1 to 3 s after noise onset. Baseline was estimated as the median value of the pre-noise period. Recovery time was defined as the interval from the trough to the first time point at which the signal reached 80% of the baseline level relative to the trough; to ensure robustness, this criterion was required to be satisfied for at least five consecutive samples. For cross-animal comparisons, values were normalized within each animal by dividing by the mean value across that animal’s conditions.

### Identification of vocalization response maps

Trials belonging to the same stimulus condition were averaged to obtain a condition-mean response movie for subsequent map computation, which was then used for the following computation. Vocalization-responsive maps were computed relative to the sustained pre-vocalization activity level immediately preceding vocalization onset. The sustained baseline window was defined as the final 100 ms preceding vocalization onset, and the vocalization response window was defined as the first 400 ms following vocalization onset, comprising the responses to the first two vocalizations.

For each pixel, the sustain baseline was computed as the mean fluorescence within the baseline window, and videos were baseline-corrected relative to this sustained activity. Vocalization response maps were computed from the baseline-corrected signal within the vocalization response window.

Binary response maps (Fig. 3A) were obtained by z-score thresholding the response maps (threshold = 1.5). The same procedure was applied for vocalizations presented during ongoing noise (Voc. in Noise) and for vocalizations presented in silence (Voc. in Sil.).

### Trace-based vocalization response metrics

For trace analysis, fluorescence signals were averaged across pixels within corresponding response maps and across trials of the same stimulus condition. To enable direct comparison between conditions, responses were extracted from a shared vocalization-responsive region (see above).

The onset response was quantified as the mean fluorescence within the first 400 ms following vocalization onset. Vocalization adaptation was quantified from the decay of the response trace during the vocalization period. Specifically, a first-order linear fit was applied to the response trace from 100 ms after vocalization onset to the end of the vocalization sequence (100-1900 ms), and the fitted slope was used as a measure of adaptation strength, with more negative values indicating stronger attenuation. To facilitate comparison between conditions, slopes were normalized by the mean slope across paired conditions within each animal.

The offset response was quantified as the rise in fluorescence following stimulus termination. The local trough was identified within the first 80 ms after stimulus offset, and the subsequent response peak was identified within 300 ms after stimulus offset. Offset-response amplitude was calculated as the difference between the mean fluorescence within ±20 ms of the peak and the mean fluorescence within ±20 ms of the preceding trough.

### Spatial separation between response areas

The spatial separation between vocalization-responsive areas was quantified using the distance between their centers of mass. For each animal and pre-vocalization duration, binary response maps were obtained separately for vocalizations presented during ongoing noise and for vocalizations presented in silence. The center of mass of each binary response map was computed from the spatial coordinates of all pixels included in the thresholded response area. Spatial separation was then defined as the Euclidean distance between the two centers of mass and converted from pixels to millimeters using the imaging scale. Distances were compared across pre-vocalization durations to assess whether the relative position of the noise-and silence-responsive regions changed with increasing exposure to the preceding noise.

### Overlap of response regions with auditory cortical fields

Spatial overlap between functional response regions and anatomical auditory areas was quantified using the Jaccard index:

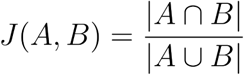

where A represents the response region and B represents the cortical field mask.

### Sorting neural response

For visualization of pixel-wise response structure, neural activity was sorted using Rastermap (Stringer et al., 2025). For each example ROI shown in Fig. 4B, a pixel-by-time response matrix was constructed from baseline-corrected widefield fluorescence traces, with rows corresponding to pixels and columns corresponding to time points. Rastermap was applied to this matrix to obtain a one-dimensional ordering of pixels based on similarity in their temporal activity patterns. The response matrix was then reordered according to this sorting and displayed as a raster plot. Rastermap sorting was used only for visualization of response structure and was not used for SVM training, testing, or statistical analyses.

### Support vector machine decoding

Neural activity was used to decode stimulus-related information using a support vector machine (SVM) classifier. For each trial, neural responses were represented as a matrix *X* ɛ R*^T×N^*, where *T* denotes the number of time frames and *N* the number of pixels within the selected region. Each time frame was treated as one sample for classification. Labels *Y* ɛ R*^T^*^×1^ were assigned according to stimulus periods. For decoding between noise-sustain and vocalization-onset periods, frames from a 1 s window preceding vocalization onset were labeled as noise-sustain, whereas frames from the 10×100 ms vocalization epochs within the vocalization sequence, with a total duration of 1 s, were labeled as vocalization-onset. A single SVM model was trained using pooled training data from all trials with pre-vocalization time ≥ 3 s to ensure stable pre-vocalization responses; both vocalization-in-noise and vocalization-in-silence trials were included. Within each trial, the noise-sustain and vocalization-onset periods were each divided into 10×100 ms bins. 70% of bins from each category were randomly selected for training and the remaining 30% were used for testing, reducing the possibility that temporally adjacent frames from being divided between the training and test sets. The classifier was implemented using the MATLAB function *fitcsvm*, with a fourth-order polynomial kernel, a box constraint of 1, and the ISDA optimization solver. Predictions were obtained using the trained model, and classification accuracy was evaluated on held-out test bins. For condition-specific analyses, test bins were grouped according to stimulus parameters, and results were summarized across animals.

### Visualization of SVM decision values

To visualize classifier separation (Fig. 4C), neural activity from a single trial was projected onto the first two principal components for display purposes only. The SVM decision value for each frame was obtained from the prediction scores of the trained model and plotted as the third dimension. Thus, the x-and y-axes represent a low-dimensional visualization of neural activity, while the z-axis reflects classifier output. The plane corresponding to zero decision value indicates the classifier decision boundary.

### Visualization of population activity using t-SNE

To visualize the organization of cortical population activity across stimulus contexts and temporal segments, t-distributed stochastic neighbor embedding (t-SNE) was applied to widefield response patterns from all pixels within the craniotomy mask. Trials with identical stimulus parameters were first averaged to obtain a condition-mean response, reducing trial-specific variability. From each condition-mean response, individual imaging frames were extracted from three temporal segments: the 1 s pre-vocalization sustain period, the 10×100 ms vocalization-onset periods within the vocalization sequence, and the nine inter-vocalization offset periods. This procedure was applied separately to vocalization-in-noise and vocalization-in-silence trials, yielding six response categories.

Each frame was represented as a vector of widefield activity values across all pixels within the craniotomy mask. For each pixel, activity values were z-scored across all included frames before dimensionality reduction. t-SNE was then applied with a perplexity of 120 to embed the high-dimensional population activity patterns into a three-dimensional space for visualization. Points in the resulting embedding represent individual frames from condition-averaged responses and were colored according to stimulus context and temporal segment. This analysis was used for qualitative visualization of representational structure and was not used to quantify classifier performance.

### Generalization across temporal segments of vocalization sequences

To assess whether decoding generalized across different temporal segments of the vocalization sequence, the classifier was trained to distinguish pre-vocalization sustain periods from vocalization-onset periods. Inter-vocalization offset periods were not included during training. Testing was performed separately on pre-vocalization sustain, vocalization-onset, and inter-vocalization offset periods. For this analysis, the classifier input consisted of neural activity from all pixels within the craniotomy mask, as for all other analyses.

Decoder output was expressed as the probability of assignment to the vocalization class derived from classification accuracy. For vocalization-onset periods, this value was defined directly as classification accuracy. For the two noise-only periods, it was computed as the inverse (1 - Accuracy), such that higher values consistently reflected stronger assignment to the vocalization-onset class across all temporal segments.

For each temporal segment, class sizes were balanced by random subsampling prior to evaluation. Chance-level output was estimated by assigning random balanced labels within each segment and recomputing the corresponding accuracy-derived measure. Analyses were performed separately for vocalization-in-noise and vocalization-in-silence trials and separately across animals.

### Decoding across pre-vocalization time and cortical areas

To assess how decoding performance depends on stimulus history and cortical region, classification accuracy was evaluated as a function of pre-vocalization time using neural activity from either primary auditory cortex (represented by an ROI centered on A1) or a vocalization-responsive region (represented by an ROI centered on the responsive region). The vocalization-responsive region was defined based on responses to vocalizations embedded in noise stimuli. For each region, a circular ROI with a radius of 200 μm was centered on the center of mass of the corresponding defined region, ensuring matched numbers of pixels and therefore matched input dimensionality for SVM analysis. A single SVM model, as described above, was trained to distinguish noise-sustain from vocalization-onset periods using pooled training data from all trials meeting the pre-vocalization time criterion. Classification accuracy was then evaluated on the held-out test data and summarized separately according to pre-vocalization time and cortical region.

### Decoding across structured noise statistics

To examine the influence of noise statistics on decoding performance, classification accuracy was evaluated across conditions defined by WFV and CFC. The same SVM model was used, and testing was performed on held-out data grouped according to the corresponding noise condition. Accuracy was computed separately for each combination of WFV and CFC. For this analysis, the classifier input consisted of neural activity from all pixels within the craniotomy mask. Class balancing and evaluation procedures were identical to those described above.

### Stimulus reconstruction from neural activity

Acoustic stimuli were converted to cochleogram representations using a gammatone filterbank spanning 2–64 kHz with logarithmically spaced frequency channels. Cochleograms were sampled at a temporal resolution of 100 Hz. Neural responses consisted of fluorescence signals from all pixels within the craniotomy mask. To account for temporal relationships between neural activity and stimulus features, a lagged design matrix was constructed by concatenating neural activity across temporal lags from −0.2 s to 0.3 s relative to each cochleogram frame. Stimulus reconstruction was performed using linear ridge regression. Let *X* denote the lagged neural response matrix and *S* the cochleogram representation of the stimulus. Reconstruction weights were estimated using ridge regression

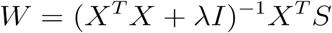

where λ is the ridge regularization parameter. It was selected empirically by comparing reconstruction performance across a range of candidate values on held-out data; λ=10000 was used for all subsequent analyses. The reconstructed cochleogram was then obtained as

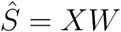

Model performance was evaluated using held-out data. Reconstruction accuracy was quantified as the Pearson correlation between reconstructed and ground-truth cochleogram signals for each frequency channel. Correlations were computed independently for each channel and then averaged across channels to obtain a single reconstruction score per trial. For population analyses, reconstruction performance was computed separately for each animal and summarized across animals.

### Area-based reconstruction and controls

To compare reconstruction across cortical regions while controlling for population dimensionality, pixels were randomly sampled from combined primary auditory areas (A1, AAF and A2), combined secondary auditory areas (DP, DM and TeA subdivisions), or across all anatomically defined auditory areas. Within each animal, the same number of pixels was used for the all-area, primary and secondary conditions, with the available number limited by the smaller cortical subdivision. Random spatial sampling was repeated and reconstruction performance was averaged across train/test splits and spatial samples to obtain one value per animal. As a control for whether additional spatial coverage improved reconstruction independently of regional identity, the all-area analysis was additionally repeated while allowing a larger number of pixels.

### Canonical correlation analysis and information estimation

CCA was performed separately for primary and secondary auditory cortex using an identical number of neural dimensions across regions and animals. 50 pixels were randomly sampled from each cortical group in every animal, and pixel sampling was repeated 10 times. Neural activity and cochleogram features from the analyzed time windows were concatenated across trials and z-scored across time before CCA. CCA identifies pairs of weighted neural and stimulus features whose temporal expressions are maximally correlated, with successive canonical components describing mutually uncorrelated relationships of decreasing strength. A chance baseline was generated by randomly permuting neural–stimulus temporal correspondence; 10 independent shuffles were performed for each pixel sample.

CCA-derived information was estimated from the canonical correlations as:

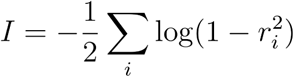

where *r_i_* denotes the canonical correlation coefficient of the *i*-th canonical dimension, and the summation is performed over all canonical dimensions. CCA spectra and information estimates were first averaged across repeated pixel samples to obtain one value per animal, and group results are reported as mean ± SEM across animals (N = 7).

### Dimensionality analysis with PCA

PCA was fitted to neural responses from the training data only, and both training and held-out responses were projected onto the resulting PCA basis. Reconstruction was then repeated using the first 1–30 principal components while keeping the underlying spatial sample and train/test partitions fixed across dimensionalities. PCA scores were not independently variance-normalized before ridge regression, thereby preserving the variance hierarchy defined by PCA. Reconstruction performance was first averaged across held-out samples within each train/test split, then across repeated splits and spatial samples to obtain one curve per animal. Group results are shown as mean ± SEM across animals (N = 7). Reconstruction using the corresponding full-dimensional neural population is shown as a reference.

## Statistical analysis

All statistical analyses were performed using MATLAB (R2022b; The MathWorks). Unless otherwise stated, data are reported as mean ± SEM across animals (N = 7). For visualization, individual data points represent animal-level averages.

Unless otherwise stated, comparisons between two conditions within the same animals were assessed using paired nonparametric tests (Wilcoxon signed-rank test, WSRT), appropriate for small sample sizes and non-normal distributions.

For experiments involving multiple conditions and repeated measurements within animals, statistical effects were assessed using linear mixed-effects models (fitlme, MATLAB). In these models, experimental factors (e.g., pre-vocalization time, variance, cross-frequency correlation, frequency, or cortical area) were treated as fixed effects, and animal identity was included as a random intercept to account for repeated measures. Interaction terms between factors were included where appropriate. Effect sizes are reported as condition means ± SEM across animals.

## Code and Data availability

The data underlying this study, as well as the data analysis files that were used in the processing of the data and creation of the figures have been deposited in the Radboud Data Repository collection (doi.org/10.34973/bnpg-kd11).

## Acknowledgements

We would like to thank Gesa Berretz, Sharon Goldewijk, Kiki Nabers, Kim van Berkel, and Samuel Varga for helping with surgery and data collection. We would like to thank Daniel Polley and Ross Williamson for discussions on setup design and Roberta Müller for assistance with the illustrations of experimental setups. BE acknowledges funding from a VIDI grant (016.VIDI.189.052) and ZC from an internal grant at the Donders Center for Neuroscience. BE also acknowledges valuable discussions at the Kavli Conference on Statistical Learning in the Brain at UCSB, supported by NSF Grant No. PHY-1748958 and the Gordon and Betty Moore Foundation Grant No. 2919.02 to the Kavli Institute for Theoretical Physics (KITP).

## Author Contributions

B.E. acquired funding, conceived and designed the experiments, managed and supervised the project, performed surgeries, and contributed to manuscript preparation. Z.C. co-developed the data-analysis pipeline, performed the primary analyses, generated the figures, assisted with surgeries, and wrote the manuscript. J.P. developed and performed the surgeries and imaging experiments and contributed to the experimental pipeline and co-development of data-analysis pipeline. S.A. contributed to stimulus-generation and analysis software and to improvements of the analysis pipeline. All authors reviewed and approved the final manuscript.

